# 3D Visualization of Macromolecule Synthesis

**DOI:** 10.1101/2020.06.24.169300

**Authors:** Timothy J. Duerr, Ester Comellas, Eun Kyung Jeon, Johanna E. Farkas, Marylou Joetzjer, Julien Garnier, Sandra J. Shefelbine, James R. Monaghan

## Abstract

Measuring nascent macromolecular synthesis *in vivo* is key to understanding how cells and tissues progress through development and respond to external cues. Here, we perform *in vivo* injection of alkyne- or azide-modified analogs of thymidine, uridine, methionine, and glucosamine to label nascent synthesis of DNA, RNA, protein, and glycosylation. Three-dimensional volumetric imaging of nascent macromolecule synthesis was performed in axolotl salamander tissue using whole mount click chemistry-based fluorescent staining followed by light sheet fluorescent microscopy. We also developed an image processing pipeline for segmentation and classification of morphological regions of interest and individual cells, and we apply this pipeline to the regenerating humerus. We demonstrate our approach is sensitive to biological perturbations by measuring changes in DNA synthesis after limb denervation. This method provides a powerful means to quantitatively interrogate macromolecule synthesis in heterogenous tissues at the organ, cellular, and molecular levels of organization.

## Introduction

The measurement of nascent DNA, RNA, and protein synthesis in animals provides critical information on the state of cells (dividing, growing, dying) in relation to their surrounding cells. Traditionally, radiolabeled or brominated nucleosides or amino acids are introduced to live animals, where they are incorporated into macromolecules during DNA synthesis^1,2^, trancription^3,4^, and translation^5^. Performing such approaches have allowed the characterization of cells actively undergoing macromolecular synthesis as well as quantification of synthesis rates, which facilitates the study of cell behavior during tissue remodeling, proliferation, stress, or disease ^6-9^.

In the past decade, bio-orthogonal macromolecule precursor analogs including 5-ethynyl-2′-deoxyuridine (EdU), 5-ethynyl-uridine (5-EU), and L-azidohomoalanine (AHA) have become commercially available. After injection of macromolecule precursor analogs into animals, precursors can later be detected in nascent DNA, RNA, and protein macromolecules with fluorescently labeled azides or alkynes through highly selective copper catalyzed azide-alkyne cycloaddition (“click”) chemistry^10-12^. These powerful new analogs provide an alternative to the use of dangerous isotopes and the challenges associated with brominated precursors such as the requirement of large secondary antibodies (∼150kd) and harsh tissue retrieval methods that limit their use in whole tissue samples.

Advantages of the click-chemistry labeling approach include the inert nature of macromolecule precursor analogs that have minimal impact on the animal, the small size of fluorescently labeled alkynes and azides, and the high selectivity of click-chemistry. These advantages have enabled whole mount fluorescent labeling of DNA synthesis^11^, RNA transcription^13^, protein translation^14^, and glycans^15,16^ in animals. These pioneering proof of principle experiments have demonstrated that imaging macromolecular synthesis is possible, but the fact that most model organisms are large, optically opaque, and consist of heterogenous tissues has made it a challenge to image biological phenomena in deep tissues. Challenges such as photon penetration, differences in refractive indices among different cellular components, light-induced photodamage, and background fluorescence have limited the use of whole mount imaging of macromolecular synthesis.

Advances in light sheet fluorescence microscopes (LSFMs) have recently enabled the imaging of large biological specimens from millimeters to several centimeters in size^17^. LSFMs have been utilized for volumetric imaging of many varieties including visualization of mRNA in whole mount fluorescence *in situ* hybridization experiments^18^ and *in vivo* interrogation of deep tissue dynamics in transgenic reporter animals^19^ among others^17^. Clearing methods like CLARITY^20^, CUBIC^21^, and 3DISCO^22^ have further enabled volumetric imaging by decreasing refractive index mismatches and tissue clearing to advance large specimen imaging even further. Together, the rise of three-dimensional imaging, new staining techniques, and tissue clearing has demanded new means for cell counting, segmentation, and fluorescence quantification.

Here, we present a click-chemistry based method to visualize DNA synthesis, transcription, translation, and protein glycosylation in whole mount samples using LSFM (Figure 1A). We demonstrate the utility of this technique by imaging macromolecular synthesis in the regenerating axolotl salamander limb. Following limb amputation, the axolotl regenerates its limb by generating a mass of proliferating cells at the limb stump called a blastema^23^. The blastema is an ideal environment to test our method because it is an accessible, heterogenous tissue that increases DNA synthesis, transcriptional output, and translation rates compared to uninjured tissue. Furthermore, DNA, RNA, and protein synthesis decrease after denervation of the regenerating limb^24,25^.

**Figure 1.**
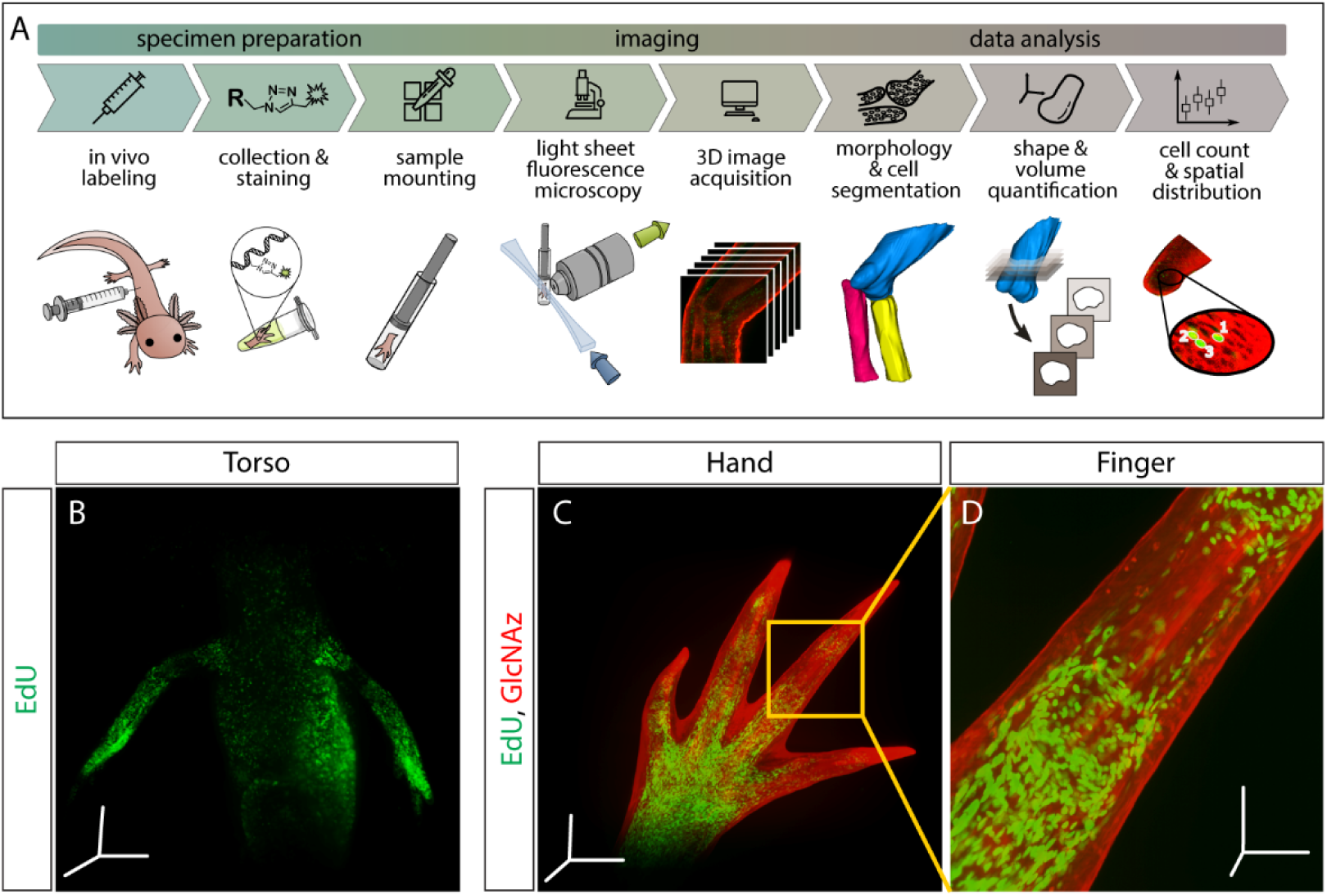
Outline of staining/analysis pipeline and exemplary images. (A) Overview of entire sample preparation, imaging, and data analysis pipeline. (B-D) Once macromolecules are labeled *in vivo*, synthesis can be visualized throughout the injected animal. Here, we show DNA synthesis (EdU) in the torso (B) and both DNA synthesis and protein glycosylation (GlcNAz) in the hand (C) and finger (D). Images from panels B and C were uncleared and imaged at 5X magnification. Image from panel D was also uncleared and imaged at 10X magnification. Scale bars for panels B&C= 600µm for each axis. Scale bars for panel D= 200µm for each axis.

We use the regenerating limb to demonstrate that two click-it ready precursors can be administered and subsequently visualized simultaneously in a single sample. We also show that optical resolution of images can be improved with the clearing agent 2,2’-thiodiethanol (TDE), and that this method works in a number of tissues. We outline an image analysis pipeline for three dimensional (3D) morphology segmentation, cell counting, and fluorescence quantification of stained tissues. We apply this pipeline to the regenerating humerus to demonstrate the multiscale quantitative analysis capabilities of our method. Finally, we show that our method is sensitive enough to detect and quantify changes in DNA synthesis rates in whole mount innervated and denervated regenerating limbs. Taken together, our method provides a unique approach to simultaneously interrogate cell state at the organ, cellular and molecular levels of organization.

## Materials and methods

### Animal procedures

Axolotls (*Ambystoma mexicanum*: d/d RRID Catalog #101L) were either bred in captivity at Northeastern University or purchased from the Ambystoma Genetic Stock Center at the University of Kentucky. Experiments were performed in accordance with Northeastern University Institutional Animal Care and Use Committee. Animals were grown to 4-6cm (Mean 5.3cm, SD 0.36) and 1-1.5g (Mean 1.3g, SD 0.19g) for use in all studies. For all experiments, animals were anesthetized by treatment of 0.01% benzocaine until visually immobilized. Limbs were amputated either at the distal end of the zeugopod or midway through the stylopod, and bones were trimmed below the amputation plane to allow for uniform growth. At the date of collection, animals were reanaesthetized and injected with either EdU (8.0ng/g animal), 5-EU (270.0µg/g animal), AHA (180.59µg/g animal), or N-azidoacetylglucosamine-tetraacylated (GlcNAz) (4.3µg/g animal) alone or simultaneously with the following combinations: EdU/AHA, EdU/GlcNAz, 5-EU/AHA, 5-EU/GlcNAz. All monomer analogs were purchased from www.clickchemistrytools.com and resuspended in DMSO at the following concentrations: EdU-300mM, 5-EU-100mM, AHA-100mM, GlcNAz-100mM. Stocks were further diluted in 1X phosphate buffered saline (PBS) for injection. Three hours after injection, limbs were collected from the upper stylopod and fixed in 4% paraformaldehyde (PFA) (diluted in 1X PBS) at 4°C overnight. If limbs were denervated, the nerve supply was severed at the brachial plexus 24 hours prior to tissue collection.

### Whole mount click-it protocol

Following fixation in 4% PFA, samples were washed three times with 1X PBS at room temperature (∼23°C) for 5 minutes. Samples were dehydrated in an increasing methanol series at room temperature starting with 25% methanol (diluted in 1X PBS), 50% methanol, 75% methanol, and 100% methanol for 5 minutes at each step. Samples could then be stored in 100% methanol indefinitely at −20°C. For staining, samples were rehydrated in a decreasing methanol series starting with 75% methanol (diluted in 1X PBS), 50% methanol, 25% methanol, and finally placed in 100% 1X PBS for 5 minutes at each step. Samples were then washed 3 times with 1X PBST (1X PBS with 1% Triton) for 5 minutes at room temperature. To aid in clearing, samples were washed in 0.5% trypsin (diluted in 1X PBST) for 30-90 minutes on a rocker at room temperature, or until sample appeared translucent. Samples were washed three times at room temperature for 5 minutes with deionized water, then washed in 100% acetone for 20 minutes at −20°C and washed with deionized water again for 10 minutes. Samples were washed in 1X PBST three times at room temperature for 5 minutes prior to applying click-it cocktail for 30 minutes at room temperature. The click-it cocktail was made in 1X TRIS buffered saline as follows: 50µL 1M sodium ascorbate (100mM final), 20µL 100mM CuSO_4_ (4mM final), and 2µL 500µM azide- or alkyne-modified Alexa Flour (2µM final), combined in order as listed. After the first round of staining, samples were washed at room temperature 6 times for 30 minutes with rocking. For double-labelling, samples were again placed in the click-it cocktail with a different fluorescent dye to stain for the second analog at room temperature overnight. Both rounds of staining were conducted in the dark to prevent photodegradation of fluorescent molecules. For staining with DAPI, samples were washed 3 times for 5 minutes with 1X PBS, then placed in 277mM DAPI for 4 days at room temperature. Samples were washed 3 times for 20 minutes with 1X PBS and left in 1X PBS at 4°C for short term storage prior to imaging with LSFM. Whole mount samples were cleared with 67% TDE (diluted in deionized water) overnight at room temperature in the dark.

### Tissue section click-it protocol

Following fixation in 4% PFA, samples were washed three times in 1X PBS each for 5 minutes, and cryoprotected in 30% sucrose on a rocker for 1 hour or until the tissue fully sinks. Samples were removed form sucrose and briefly washed in optimal cutting temperature (OCT) compound before mounting in OCT compound and frozen at −80°C. 10µm sections were obtained using a cryostat and baked at 65°C for 15 minutes. Slides were then washed with water for 30 minutes at room temperature to remove residual OCT. Slides were washed once with 1X PBS for 5 minutes at room temperature. The click-it cocktail (same as above) was applied to the slides and incubated at room temperature in the dark for 30 minutes. If staining for a second macromolecule, slides were washed five times for 5 minutes with 1X PBS at room temperature. The samples were then stained for 30 minutes at room temperature in the dark using the above click-it cocktail with a different a fluorophore dye. Following the final click-it reaction, slides were washed once with 1X PBS at room temperature for 5 minutes, then stained with 277mM DAPI for 5 minutes at room temperature. Slides were washed again with 1X PBS for 5 minutes at room temperature and water for 5 minutes at room temperature and mounted with SlowFade™ Gold Antifade Mountant (Thermo S36936). Slides were imaged using a Zeiss LSM800 confocal microscope.

### Light sheet microscopy

All 3D images were acquired using a Zeiss light sheet Z.1 microscope paired with Zen software. Unless otherwise indicated, samples were cleared and imaged in 67% TDE. Post-processing for visualization purposes was performed with Arivis Vision4D v3.1.4 on a workstation with a 64-bit Windows Embedded Standard operating system, and an Intel(R) Xeon(R) CPU E5-2620 v3 @ 2.40GhZ (2 processors), 128GB RAM, and NVIDIA Quadro K2200 GPU. Sub-volumes were stitched together with the Tile Sorter in Arivis, using the manual projection option. Volume fusion was performed through automatic landmark registration of manually selected points for alignment. Finally, constant background intensity was corrected using the automatic functionality.

### Data analysis

All data were processed on desktop computers with the aid of Fiji^26^ and Matlab^27^. The custom Fiji scripts and Matlab codes used are available in the supplementary material. We performed all analyses on unprocessed .czi files acquired directly from the light sheet microscope.

#### Organ level

The goal of the organ level analysis was to determine gross morphology of the organ, such as area, principal axes, and maximum diameters in said axes for each cross section. Critical to this step is alignment of the images to a standard defined axis and reslicing in the transverse direction (perpendicular to the long axis of the humerus). The image stack of a regenerating axolotl elbow was imported into Fiji and aligned along the proximodistal axis of the humerus (Figure 3A). Morphological segmentation of the humerus was performed semi-automatically with the Segmentation Editor plugin (Figure 3B). The segmented surface was then exported from Fiji as a mesh via an .stl file using 3D Viewer^28^. Volumetric analysis was performed in Fiji by reslicing the aligned mask (Figure 3B) to obtain a stack of cross-sections perpendicular to the proximodistal axis, and then quantified the shape and volume with the Slice Geometry option in the BoneJ^29^ Fiji plugin. Slice Geometry calculates cross-sectional geometric properties of shapes, including area, second moment of area around the major and minor axes, and maximum chord length from these major and minor axes.

**Figure 2.**
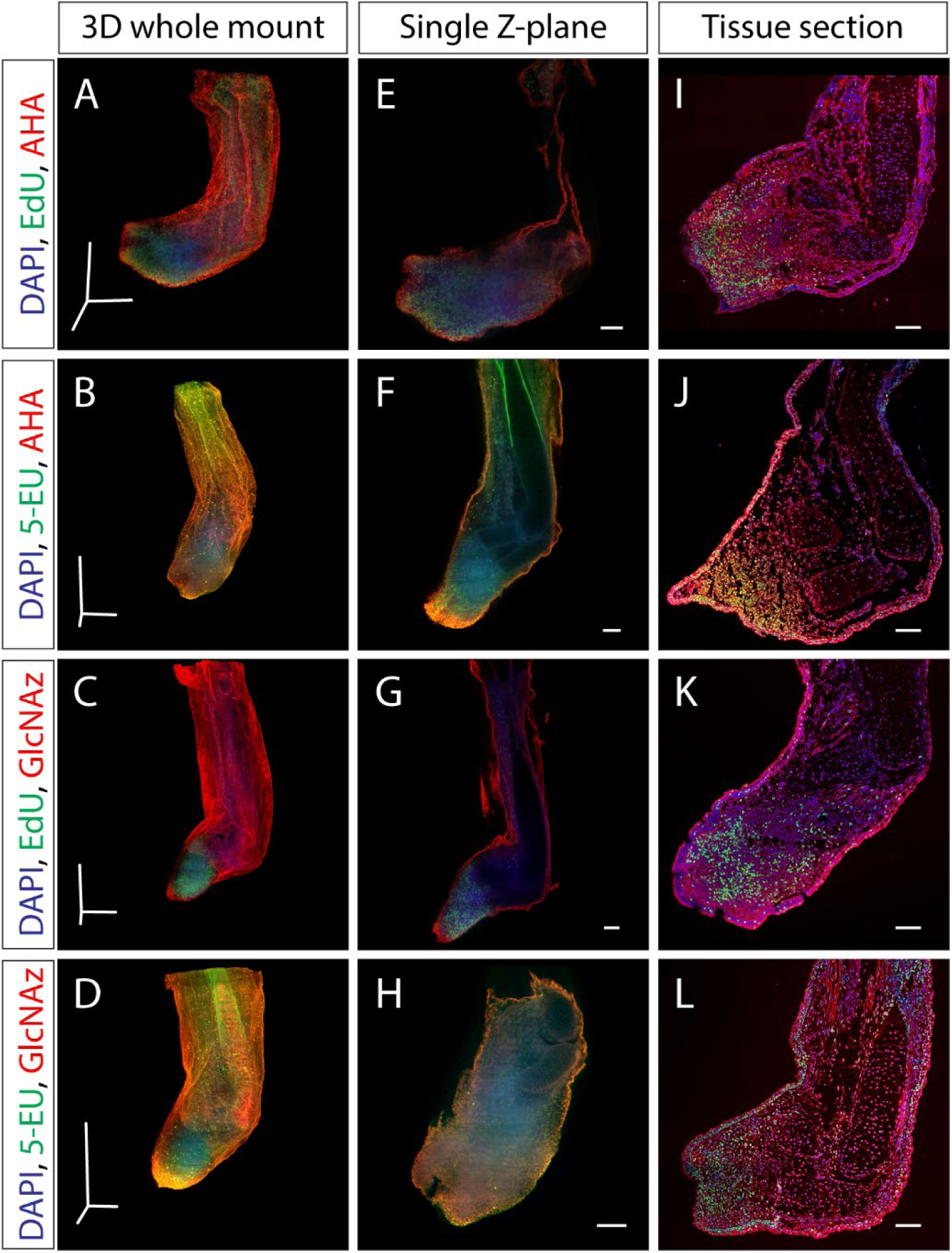
Dual staining of macromolecule synthesis in whole mount. (A-D) Stitched and fused 3D reconstruction of 13 dpa blastemas stained for multiple macromolecules obtained by LSFM. (B-H) Single Z-plane from A-D that represents the entirety of the blastema. (I-L) Tissue section from identically treated limbs as A-H showing similar macromolecule staining patterns, indicating that the whole mount staining method does not alter macromolecule synthesis staining patterns. Scale bars for panels A-D= 600µm for each axis. Scale bars for panels E-L= 200µm.

**Figure 3.**
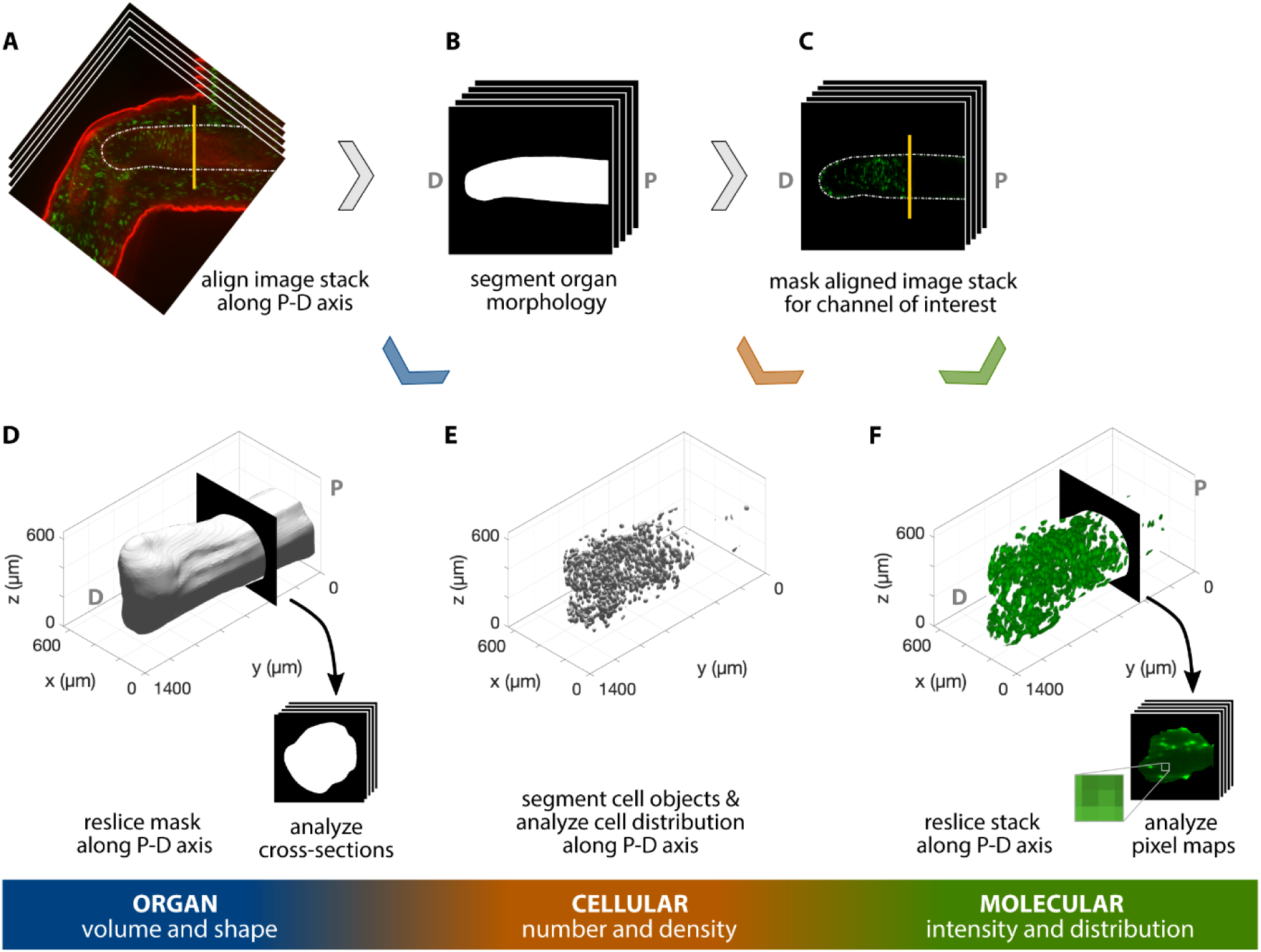
Workflow for 3D, multiscale analysis of the regenerating axolotl humerus. Multiscale analysis of a 35 dpa regenerating axolotl humerus, stained for AHA (red) and EdU (green). The humerus in the image stack was (A) aligned along the proximodistal (P-D) axis and (B) its morphology was segmented. The resulting mask was used to analyze the organ volume and shape by (D) reslicing it along the P-D axis and studying the cross-sections obtained. (C) The segmented morphology in B was used to mask the green channel for cellular- and molecular-level analyses. (E) Cells in the humerus were segmented and their spatial distribution was analyzed to obtain cellular number and density. (F) The masked image stack in C was resliced along the P-D axis and the pixel maps of the cross-sections were used to characterize the molecular intensity and distribution within the humerus. The vertical yellow line in A and C indicates the plane of amputation.

#### Cellular level

The goal of the cellular level analysis was to determine the number of proliferating cells (EdU^+^) as a function of proximodistal position along the humerus, as well as average size, shape, and orientation. The aligned image stack (Figure 3A) was cropped using the morphological segmentation (Figure 3B) as a mask with the aid of the Image Calculator in Fiji. Based on the EdU staining (Figure 3C), we segmented the nuclei of the proliferating cells with Trainable Weka Segmentation 3D^30^. We used a combination of filters available in Fiji before and after training the algorithm to improve segmentation results. Filters applied include the 3D Edge and Symmetry, Background Subtraction, 3D Fill Holes, Gaussian Blur 3D, and 3D Watershed Split. A detailed description of the sequence and parameters used is available in the supplementary materials. The 3D Objects Counter then provided a list of identified cells as well as their volume and position, among other information. In addition, the surfaces of the segmented cell nuclei were exported and processed in MeshLab, similarly to the process followed with the organ segmentation, to be visualized in Matlab (Figure 3E). The centroid coordinates and corresponding object volumes identified in Fiji were imported into Matlab for further processing and plotting. We computed the number of cell centroids within 50μm thick slices perpendicular to the proximodistal axis to obtain a density-like measure of proliferating cells. Slice thickness was selected as slightly larger than the average cell diameter to ensure cells were not sampled across more than two slices.

#### Molecular level

The goal of the molecular level analysis was to determine molecular activity, characterized by fluorescent signal, which in our study represented DNA replication (EdU). The aligned and cropped image stack used as a starting point of the cellular-level analysis (Figure 3C) was also analyzed at the molecular level. We resliced the image stack along the proximodistal axis in the organ under study (Figure 3F). The histogram and pixel intensity statistics were listed for each slice via the getRawstatistics function. These allow for further quantification of the fluorescent signal in Matlab, e.g. we calculated the mean intensity of each plane along the proximodistal axis and the histogram of the whole organ. For the example showing nerve-dependent regeneration in axolotl blastemas (Figure 6A), we compared overall pixel intensity of the left (denervated) and right (control, innervated) forelimbs of the same animal. To ensure full repeatability of our data analysis, we selected the same cubic volume in all blastemas processed: a cube with 175μm sides, centered along the proximal-distal axis and at a distance of 250μm from the distal tip (Figure 6A). The cube size was adjusted to maximize the cube volume for all blastemas processed, while ensuring that the entire cube was contained within the blastema. Histograms of each cubic volume and the ratio between the innervated and denervated limb of the same animal were computed and plotted with Matlab (Figure 6B) to quantify the changes in proliferation in innervated versus denervated limbs at different stages of regeneration. All scripts for blastema cube quantification are provided in the supplementary materials.

**Figure 4.**
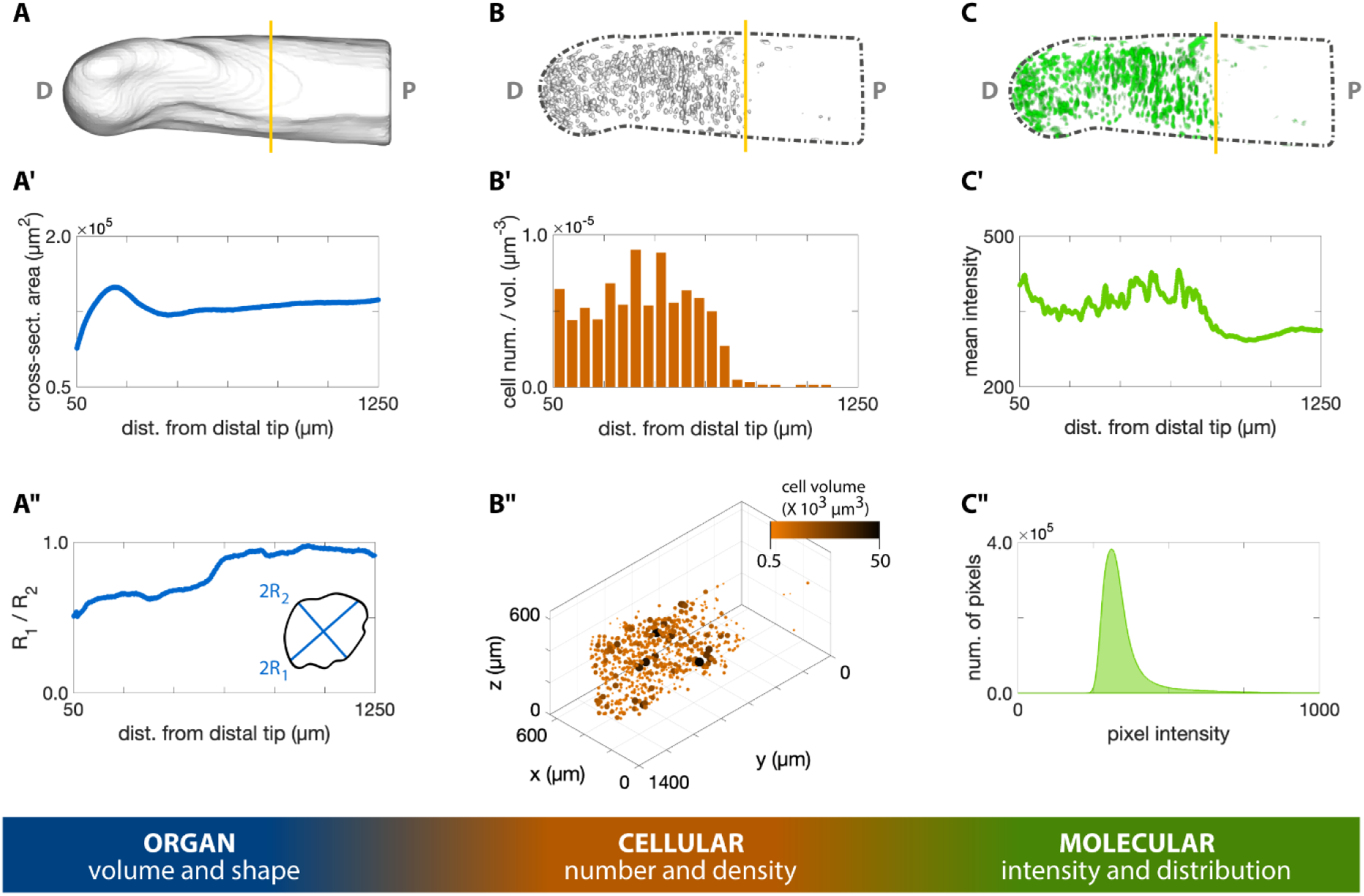
3D quantification across scales of a regenerating axolotl humerus. (A) The cross-sections of the humerus in Figure 3D were analyzed with the Fiji plugin BoneJ to quantify humerus shape and volume. (A’) The cross-sectional area along the proximodistal (P-D) axis provides a measure of volume distribution along the humerus. (A”) The ratio of the maximum chord length from the minor axis (2R_1_) with respect to the maximum chord length from the major axis (2R_2_) provides a measure of cross-sectional circularity in the humerus. Values closer to 1.0 in the proximal side indicate a more circular cross-section in this zone. (B) The Fiji plugin Trainable Weka Segmentation 3D and 3D Objects Counter were used in the cellular analysis of proliferating chondrocytes illustrated in Figure 3E. (B’) The number of EdU^+^ cells within a 50µm slice along the P-D axis was divided by the slice volume to obtain a density-like measure. (B”) The center of mass of each cell was plotted in 3D, with point size and color proportional to the segmented cell volume. (C) The molecular intensity and distribution were analyzed based on the resliced pixel intensity maps of the masked green channel in Figure 3F. (C’) Mean intensity of each slice perpendicular to the P-D axis. (C”) The histogram of the EdU staining in the humerus can provide a measure of DNA synthesis rate. The vertical yellow line in the top row images indicates the plane of amputation.

**Figure 5.**
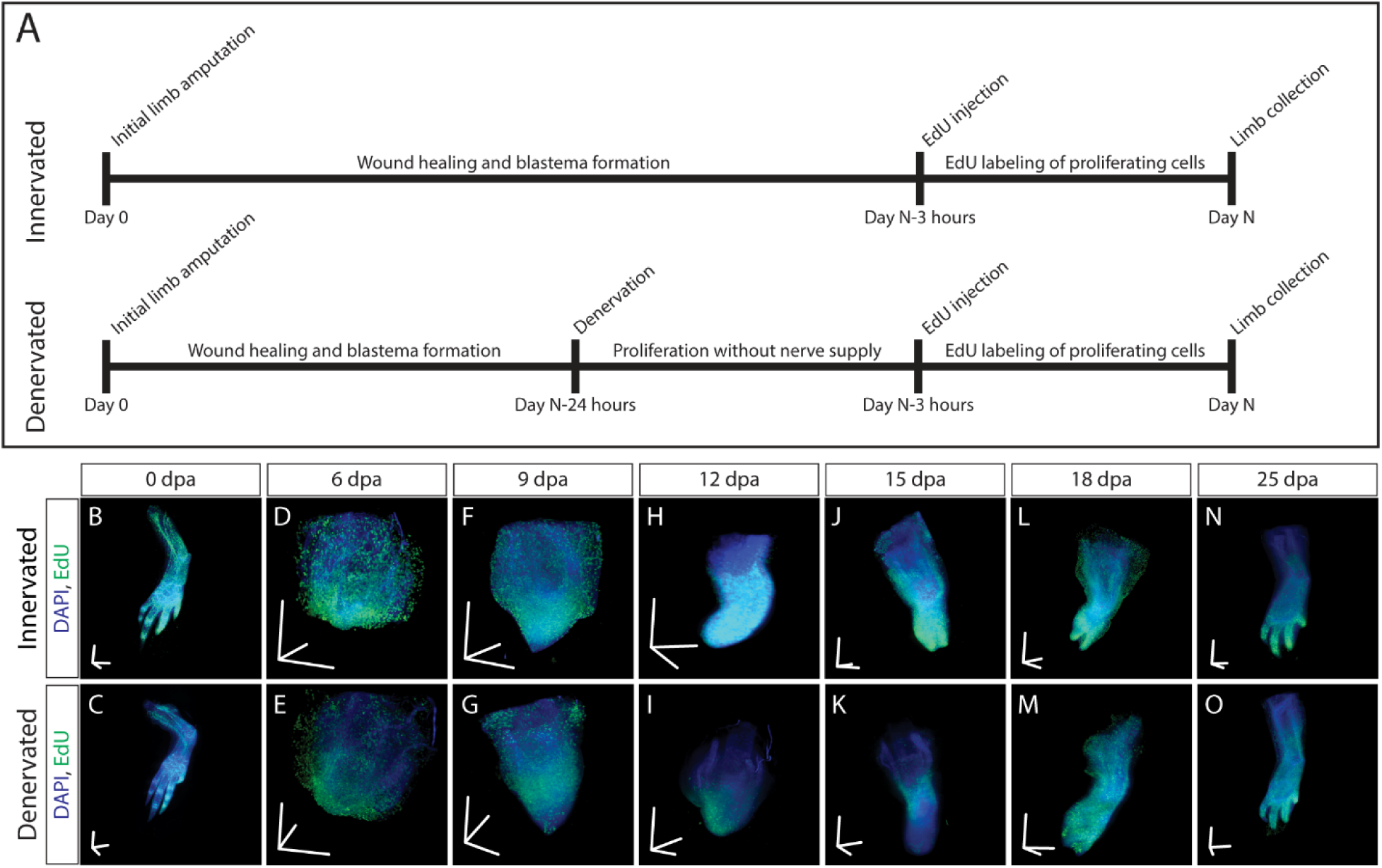
3D visualization of DNA synthesis in innervated/denervated regenerating limbs. (A) Schematic of experimental design used to obtain samples from B-O. (B-O) Time course of regeneration in innervated and 24 hour denervated limbs at 0, 6, 9, 12, 15, 18, and 25 dpa. Scale bars for panels B-O=600µm for each axis.

**Figure 6.**
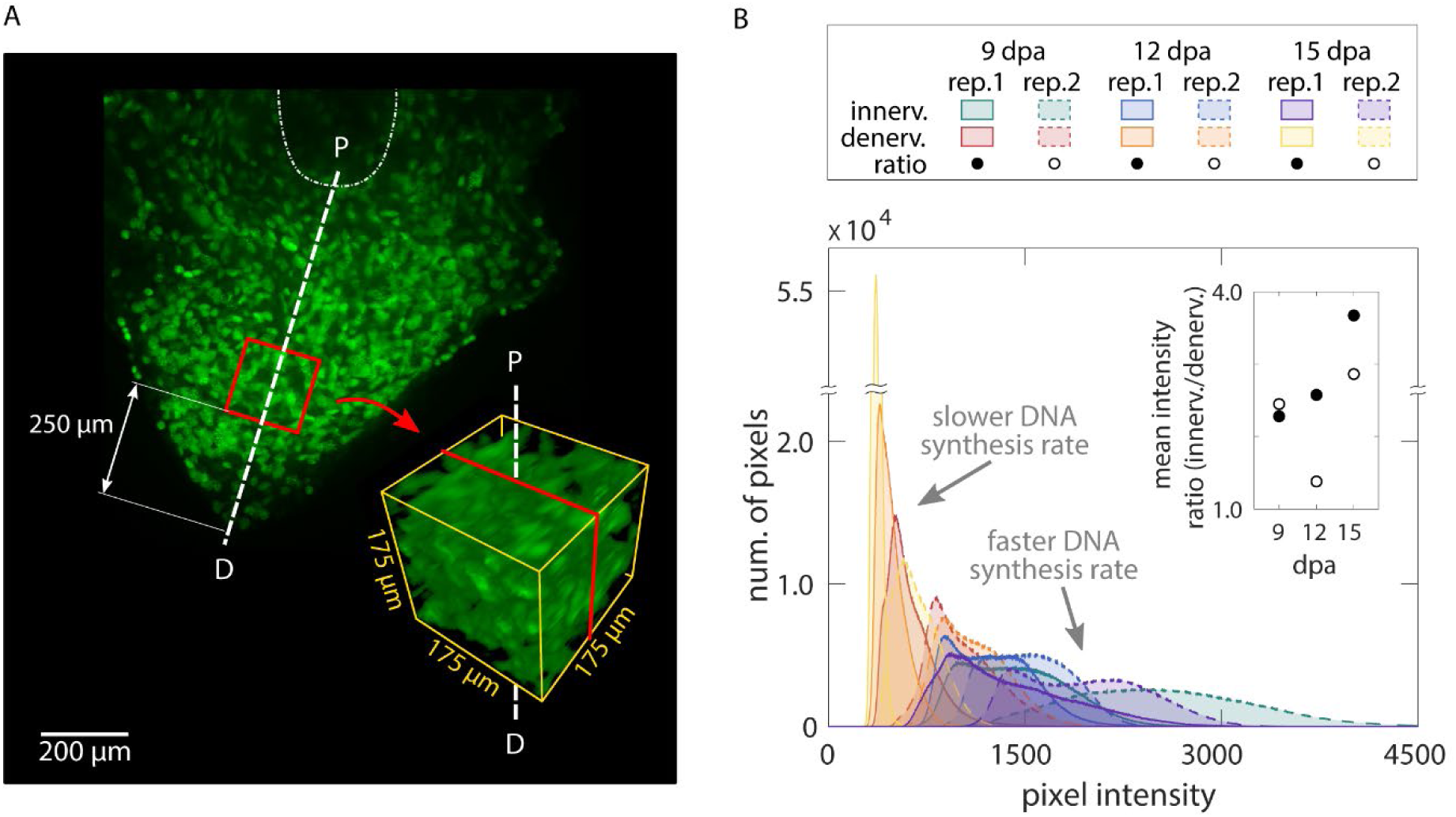
3D quantification of DNA synthesis in innervated/denervated regenerating limbs. (A) A cube with sides of 175µm was cropped along the proximodistal axis 250µm from the distal tip of each blastema. (B) The overall histogram of the cubes confirms that innervated blastemas have faster DNA synthesis rates than their denervated counterparts. The difference is quantified by computing the ratio between the mean pixel intensity of the innervated versus denervated blastema cube of a same animal (inset plot).

## Results and discussion

### Whole mount, click-it based visualization of macromolecule synthesis

To visualize macromolecule synthesis, we injected click-it compatible monomer analogs (Table 1) intraperitonially three hours prior to sample collection. During this time, analogs metabolically incorporated into nascently synthesized macromolecules, resulting in *in vivo* labeling of macromolecule synthesis. These labeled macromolecules contain either azide- or alkyne-modified monomers that can be detected with click-it compatible fluorescent molecules, enabling imaging of nascent macromolecules in whole mount tissues with LSFM (Figure 1). To demonstrate the multiplexing capabilities of the approach, modified monomer analogs with disparate functional groups (EdU/AHA, 5-EU/AHA, EdU/GlcNAz, 5-EU/GlcNAz) were co-injected and visualized with LSFM (Figure 2A-D, Supplementary videos 1-2). Whole mount samples were comparable to 2D longitudinal tissue sections of the same stains (Figure 2I-L), showing that our method generates similar results in both whole mount and tissue sections (Supplementary Figure 1). We demonstrate the specificity of N-Acetylglucosamine (GlcNAc) incorporation by showing that a GlcNAz-specific antibody pretreated on tissue sections collected from GlcNAz injected animals prevented the subsequent click-it reaction (Supplementary Figure 2).

**Table 1.** Monomer analogs used to demonstrate whole mount visualization method.

| **Name** | **Macromolecule analog** | **Biological process** | **Click-it modification** | **Reference** |
| --- | --- | --- | --- | --- |
| 5-ethynyl-2'-deoxyuridine (EdU) | Thymidine | DNA synthesis | Alkyne | Salic & Mitchison 2008 |
| 5-Ethynyl Uridine (5-EU) | Uracil | Transcription | Alkyne | Jao & Salic 2008 |
| L-Azidohomoalanine (AHA) | Methionine | Translation | Azide | Wang et al. 2017 |
| Azide-modified glucosamine (GlcNAz) | Glucoseamine | Protein glycosylation | Azide | Laughlin et al. 2006 |

An advantage of our method is that staining whole mount tissues eliminates the need for sectioning, reducing the potential inconsistences that arise as a result of the sectioning process (uneven tissue, different cutting planes, etc.). Additionally, traditional methods of obtaining 3D images of thick tissues with confocal microscopy are impractically slow when imaging hundreds of images in a single stack. LSFM allows for more rapid imaging of whole samples, requiring only minutes to image each sample. However, several considerations exist when imaging whole tissue in 3D with LSFM. Stain penetrates slower in whole 3D tissues compared to 10-20µm thick tissue sections, requiring longer staining times. An advantage of our method is that the click reaction requires molecularly diminutive reagents that readily pass through cell and nuclear membranes, ensuring stain penetration in the center of dense tissues. Different refractive indices between disparate cellular components and the imaging media cause light to scatter, which can reduce the resolution and brightness of 3D images^31^. To improve image resolution and light penetration, overnight refractive index matching with 67% 2,2’-thiodiethanol (TDE) can sufficiently clear axolotl limbs for imaging with LSFM. This clearing method rapidly and effectively improves the signal to noise ratio of stained samples compared to imaging in PBS (Supplementary Figure 3). With careful attention to these challenges, our method provides a means to obtain high quality 3D images from tissues 1mm in depth in less than 10 minutes per sample (Figure 2A-H) with clear, consistent staining.

### 3D, multiscale quantitative analysis of the regenerating humerus

To demonstrate that our whole mount click-it method can obtain quantifiable data on the organ, cellular, and molecular levels of organization, we applied our technique to the regenerating humerus. After regenerating for 35 days, axolotls with mid-humeral amputations were injected with EdU/AHA to identify cells within the humerus undergoing DNA synthesis (EdU) and protein translation (AHA). We observed EdU staining in chondrocytes distal to the amputation plane and AHA staining in the humerus perichondrium (Figure 3A). We outline a multiscale, quantitative pipeline that leverages the staining patterns of these macromolecules for analysis of 3D humerus morphology and 3D macromolecule synthesis. This workflow combines available plugins in Fiji^26^ and scripts developed in Fiji and Matlab^27^ for the data analysis process (see supplementary information for a detailed description).

For 3D organ level analysis of macromolecule synthesis, AHA staining provided an adequate outline of the humerus, allowing us to segment its 3D morphology (Figure 3A-B). We quantified organ shape by assessing cross-sectional area (Figure 4A’) and circularity (Figure 4A”) of the segmented humerus along the proximodistal axis with BoneJ^29^ (Figure 4A). These measurements of 3D organ size can be obtained with other methods such as microCT and focused ion beam scanning electron microscopy (FIB-SEM). However, microCT is unable to image every tissue due to stain limitations and uses hazardous radiation, while FIB-SEM can capture 3D surface topography but not the entire organ morphology. Our method has the capability to fully image the 3D structure of entire organs (size permitting) without the need for radiation while simultaneously capturing information from both the cellular and molecular levels of organization.

On the cellular level, we segmented the 3D morphology of proliferating chondrocytes based on EdU staining using the Trainable Weka Segmentation 3D plugin^30^ (Figure 4B”). With this segmentation, we identify highly condensed regions of cells undergoing rapid rates of DNA synthesis (Figure 4B). From these data, we observe cells synthesizing DNA most abundantly distal to the plane of amputation, as expected for dividing chondrocytes. Traditionally, cell quantification as such is conducted on 2D tissue sections. In heterogenous tissues, however, cells are distributed non-uniformly; 2D sampling may not accurately capture the cellular distribution and cannot be used to determine cell volume or shape. Our whole mount staining method allows quantification of cells within an entire 3D tissue, resulting in a more accurate assessment of cell density within heterogenous tissues.

To demonstrate quantitative molecular analysis of 3D macromolecule synthesis, we assessed staining intensity of EdU in slices along the proximodistal axis of the regenerating humerus (Fig 4C). EdU intensity represents the rate at which a cell undergoes DNA synthesis, which is one of the first steps of cell division and proliferation. Proliferating cells rapidly synthesize DNA, enabling cells to integrate EdU into nascent DNA strands, providing ample opportunities for covalent linkage of fluorescent molecules to the DNA strand. Thus, higher pixel values within EdU^+^ cells are representative of faster DNA synthesis rates. These data show that EdU intensity is strongest in the regions distal to the amputation plane of the humerus (Figure 4C’), providing a quantitative measure to assess macromolecule synthesis on the molecular level. Within a tissue, cells synthesize macromolecules heterogeneously, reflected by different fluorescence values between cells. Thus, quantifying macromolecule synthesis based on presence/absence of signal instead of fluorescence does not account for variability in macromolecule synthesis rates among cells. To compound this issue, quantifying fluorescence in tissue sections only represents the rate of macromolecule synthesis from a fraction of cells in larger, heterogenous tissues. Our method provides a means to capture this molecular heterogeneity in 3D samples, allowing us to observe whole 3D regions in the regenerating humerus that synthesize DNA more rapidly than others, which further demonstrates the utility of our whole mount click staining method.

Taken together, these results demonstrate that our method can provide quantifiable data on the organ, cellular, and molecular levels of organization. This highlights the novelty of our method, as we have not found previous examples of multiscale analysis as outlined here. We foresee this multiscale, quantitative analysis having broad applications in the examination of dynamic cell processes in 3D, such as in cancer metabolism and mammalian neurogenesis or other fields where macromolecule synthesis is traditionally studied in tissue sections.

### 3D, molecular analysis of biological perturbations

To demonstrate that our method is sensitive enough to detect subtle changes in macromolecule synthesis in vivo, we quantified the difference in EdU intensity between limbs regenerating with and without a nerve supply. Cell proliferation in regenerating limbs is sustained by factors secreted from nerves that innervate the limb^32,33^. Therefore, we amputated both forelimbs at the mid-humerus, and the left limb was denervated at the brachial plexus one day prior to collection. At 6, 9, 12, 15, 18, and 25 days post amputation (dpa) animals were pulsed with EdU for 3 hours before collection to label proliferating blastema cells (Figure 5A). LSFM was used to image samples (Figure 5B), ensuring pixel resolution was consistent between samples. We quantified DNA synthesis in denervated limbs compared to innervated limbs by creating a 175×175×175µm cube 250µm from the distal most tip of the blastema (Figure 6A). From these results, we observed a marked decrease in blastema EdU incorporation due to denervation at 9, 12, and 15dpa (Figure 6B), demonstrating that our whole mount staining approach is capable of detecting changes in macromolecule synthesis after biological perturbations. One potential limitation of our method in the blastema is the inability to preform single cell segmentation. We predict that imaging with higher magnification may overcome this limitation but will significantly increase imaging time and file size.

## Conclusions

The work presented here provides a fast, simple pipeline for visualizing macromolecule turnover in the 3D space of whole tissues. While tissue sections were previously the standard for studying these cellular processes, a new standard in the field must be expected where dynamic processes like macromolecule synthesis are visualized in 3D to obtain a more complete understanding of how these processes occur in a larger tissue context. Few modalities of imaging exist to provide this level of analysis. Here, we outline a method to study macromolecule synthesis at the organ, cellular, and molecular levels of organization, which is important in understanding cell state and how cell state affects neighboring cells and tissues. To this end, we show that DNA synthesis, transcription, translation, and protein glycosylation increase in the entire 3D space of the regenerating blastema after limb amputation and that these processes can be visualized concurrently. We outline a multiscale pipeline for analysis and quantification of heterogeneous tissues at the organ, cellular, and molecular levels of organization. Additionally, we demonstrate that our method is sensitive to detect biological perturbations by showing a decrease in DNA synthesis in the blastema following limb denervation. We foresee our method being used to similarly readdress other classical questions with modern techniques for a more exhaustive understanding of biological processes; traditional questions within the fields of cancer biology and neurobiology may especially benefit from a technology as such. Additionally, as more click-it ready macromolecules monomer analogs are generated, our method will provide a means to study more biological processes in whole tissues.

## Acknowledgements

The authors would like to acknowledge Tyler Jensen for his early assistance in developing the whole mount staining protocol, and both Alex Lovely and Gouxin Rong for their microscopy and image analysis assistance. Images obtained from the Harvard University Center for Biological Imaging and the Northeastern University Chemical Imaging of Living Systems core. We acknowledge animal support from the Ambystoma Genetic Stock Center funded by NIH grant P40-OD019794.

## Author Contributions

SJS and JRM conceived and supervised the study. TJD, EKJ, and JEF preformed experiments and imaging. EC, MJ, and JG designed and implemented codes for multiscale data analysis. TJD and EC performed the analysis. TJD, EC, SJS, and JRM wrote the paper. All authors approved the final version of the manuscript.

---

**Supplementary figure 1.**
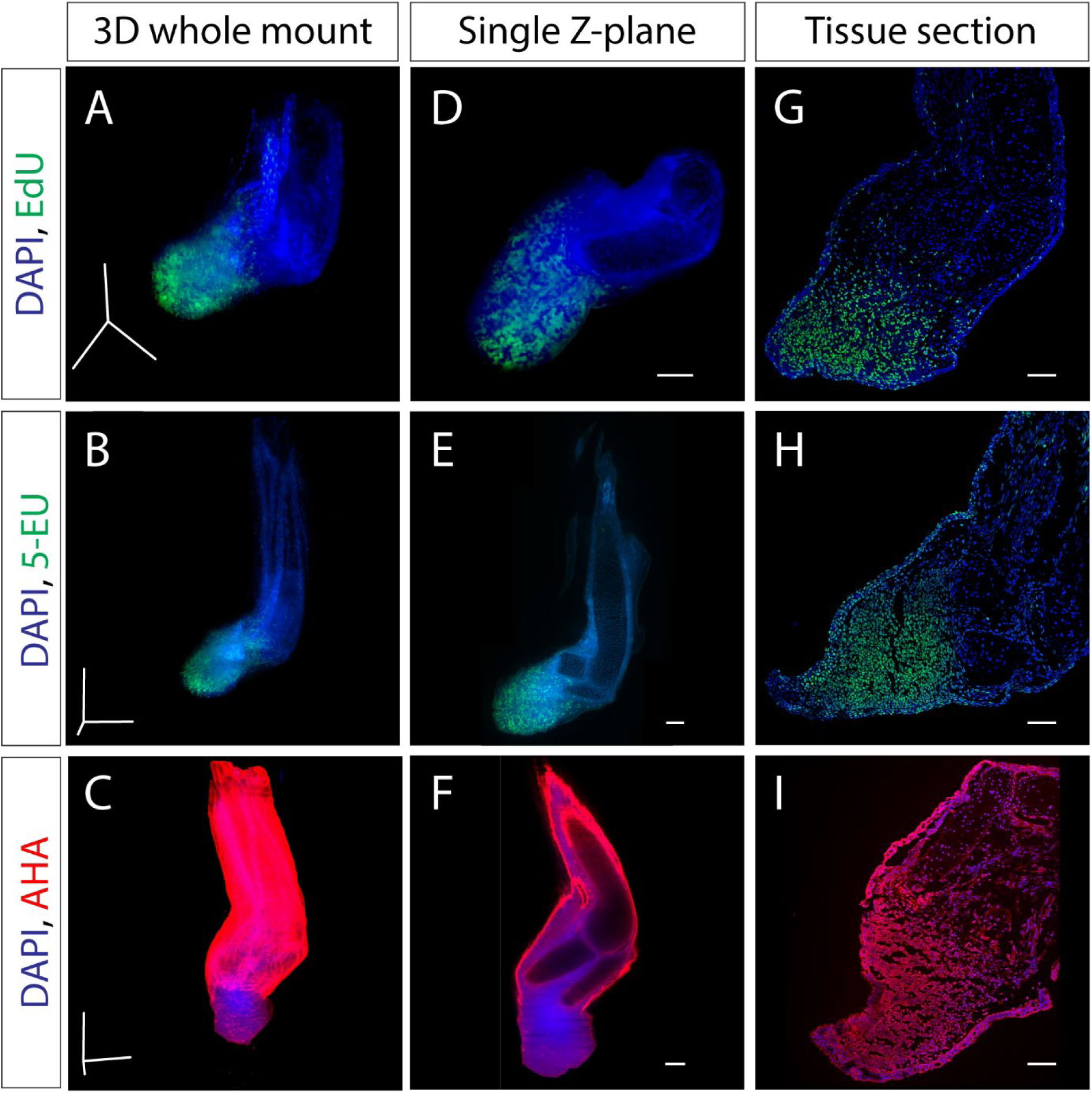
Single staining of macromolecule synthesis in whole mount. (A-C) Stitched and fused 3D reconstruction of 13 dpa blastemas stained for one macromolecule obtained by LSFM. (D-F) Single Z-plane from A-C that represents the entirety of the blastema. (G-I) Tissue section from identically treated limbs as A-F. Scale bars for panels A-C= 600µm in each axis. Scale bars for panels D-I= 200µm.

**Supplementary figure 2.**
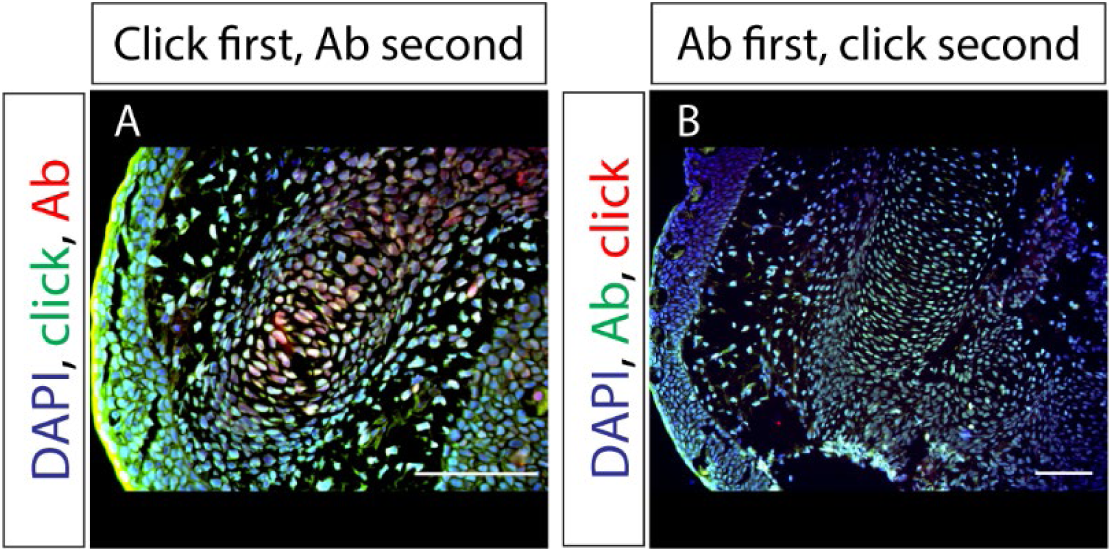
Specificity of GlcNAz staining. (A) Tissue section of a regenerating axolotl limb where the click-it cocktail (green) for GlcNAz was applied prior to staining with GlcNAc antibodies (Ab) (red). (B) Tissue section of a regenerating axolotl limb where GlcNAc antibodies (green) were applied prior to treatment with the click-it cocktail (red) for GlcNAz, demonstrating the specificity of the *in vivo* GlcNAz labeling. Scale bars=100µm.

**Supplementary figure 3.**
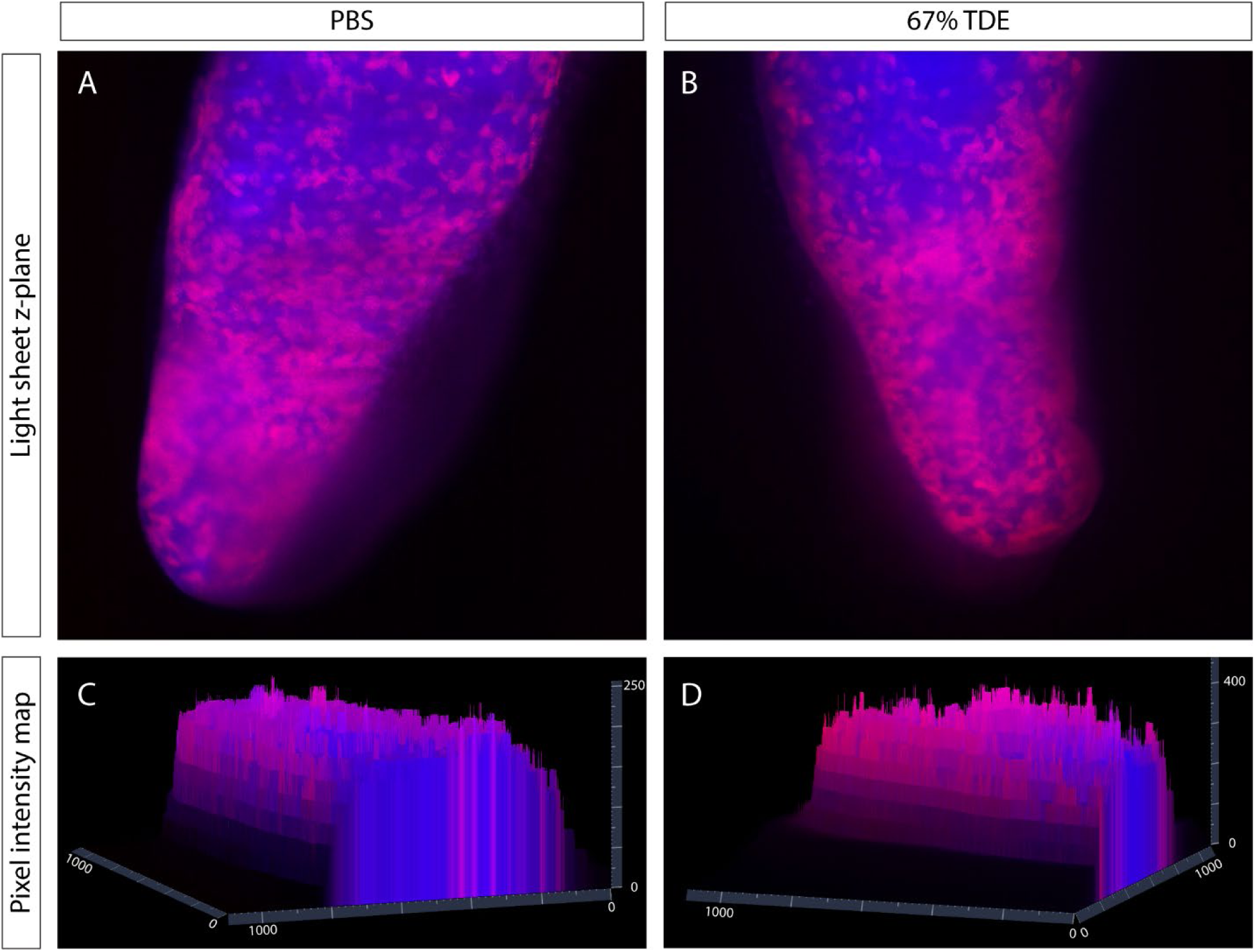
Comparison of imaging in PBS and 67%TDE. (A-B) Single Z-plane of 13 dpa blastema imaged in PBS (A) or cleared and imaged in 67% TDE (B). Red indicates EdU staining whereas blue represents DAPI staining. (C-D) Pixel intensity map of PBS imaged blastema (C) and 67% TDE imaged blastema (D). Scale bars are in units of microns.

**Supplementary video 1**

Rotating axolotl hand stained for EdU (green) and GlcNAz (red).

**Supplementary video 2**

Rotating axolotl limb stained for EdU (green), AHA (red), and DAPI (blue).

**Supplementary video 3**

Scroll through of Z-stack from supplementary video 2.

